# Dual genome-wide CRISPR knockout and CRISPR activation screens identify common mechanisms that regulate the resistance to multiple ATR inhibitors

**DOI:** 10.1101/2020.04.08.032854

**Authors:** Emily M. Schleicher, Ashna Dhoonmoon, Kristen E. Clements, Lindsey M. Jackson, Coryn L. Stump, Claudia M. Nicolae, George-Lucian Moldovan

## Abstract

The ataxia telangiectasia and Rad3-related (ATR) protein kinase is a key regulator of the cellular response to DNA damage. Due to increased amount of replication stress, cancer cells heavily rely on ATR to complete DNA replication and cell cycle progression. Thus, ATR inhibition is an emerging target in cancer therapy, with multiple ATR inhibitors currently undergoing clinical trials. Here, we describe dual genome-wide CRISPR knockout and CRISPR activation screens employed to comprehensively identify genes that regulate the cellular resistance to ATR inhibitors. Specifically, we investigated two different ATR inhibitors, namely VE822 and AZD6738, in both HeLa and MCF10A cells. We identified and validated multiple genes that alter the resistance to ATR inhibitors. Importantly, we show that the mechanisms of resistance employed by these genes are varied, and include restoring DNA replication tract progression, and prevention of ATR inhibitor-induced apoptosis. Our dual genome-wide screen findings pave the way for personalized medicine by identifying potential biomarkers for ATR inhibitor resistance.

## Introduction

Proper response to DNA damage and replication stress is critical for all organisms. Replication stress occurs upon arrest of the DNA replication machinery at sites of DNA damage, or during replication of endogenous difficult to replicate DNA sequences such as microsatellite regions^1^. The ataxia-telangiectasia-mutated (ATM) and ataxia telangiectasia and Rad3-related (ATR) kinases are in primary control of the cellular responses to replication stress and DNA damage^2^. The activation of these kinases is critical to arrest the cell cycle and allow time for proper execution of DNA replication and repair prior to cell division^3^. ATR is activated by single-stranded DNA (ssDNA) formed upon replication fork arrest. ATR activation leads to downstream phosphorylation of Chk1, resulting in stabilization of the replication fork, suppression of origin firing, and cell cycle arrest. ATM is triggered by the presence of double-stranded DNA breaks, and phosphorylates P53 and Chk2 leading to cell cycle arrest to allow for the DNA to be repaired before proceeding through the cell cycle.

Cancer cells heavily rely on the replication stress response for viable cell division^4^. Many of the current cancer treatment agents are genotoxic compounds that lead to a variety of adverse side effects for patients, as non-tumor cells in the body can also be affected. One way to avoid these side effects is to enhance the specific targeting of cancer cells, by exploiting their reliance on the replication stress response. Thus, targeting ATR has been proposed as a potential cancer therapy^5^. ATR inhibitors (ATRi) could be an option for killing cancer cells with an inherently large amount of DNA damage, due to the pivotal role of ATR in the DNA damage response^6^. Additionally, non-tumor cells in the body have little to no replication stress, and thus should not be affected by ATRi. ATR inhibitors may also be used in combination with DNA damaging agents as a therapeutic option^7,8^. Moreover, since ATR and ATM work together to manage DNA damage within the cell, ATR inhibitors have been shown to be particularly efficient in ATM/P53 deficient tumor cells^9–11^.

Since the development of ATR inhibitors, the effects of ATR/Chk1 pathway inhibition have been examined more closely. It was originally shown that inhibition of Chk1 caused a decrease in the inter-origin distance during DNA replication, accompanied by a decrease in replication fork progression^12^. Under normal conditions, Chk1 inhibits replication initiation by blocking CDC45 recruitment to the MCM2-7 complex, which is necessary for unwinding DNA at the replication fork^13^. Much like loss of Chk1, ATR inhibition has been shown to cause a decrease in replication fork speed as well as an increase in the amount of origins that are firing during DNA replication^14,15^.

AZD6738 and VE822 (M6620/Berzosertib/VX-970) are two ATR inhibitors currently under investigation in multiple clinical trials. AZD6738 is an ATP competitive, orally bioavailable ATR inhibitor^11^. AZD6738 blocks the phosphorylation of Chk1-Ser345, the downstream target of ATR. AZD6738 inhibits both ATR kinase activity and Chk1 phosphorylation at an IC_50_ of 1 nM and 74nM, respectively. Additionally, AZD6738 does not significantly inhibit other PI3K-like kinases such as ATM^11^. VE822 also blocks the phosphorylation of Chk1-Ser345^16^. VE822 and AZD6738 are currently in 11 phase I and phase II clinical trials and are being tested alone or in combination with other drugs such as Olaparib, Cisplatin, Paclitaxel, etc^17^. In addition, some of the current clinical trials focus on patients with particular tumor biomarkers such as DNA damage response pathway mutations, DNA damage, P53 mutations, or ATM deficiency^17^. Early results of these clinical trials show that both ATR inhibitors have low toxicity in patients and work best in combination with other drugs^18,19^. Specifically, VE822 was shown to be well tolerated in phase I trials with low toxicity and no dose-limiting affects^20^. Similarly, AZD6738 was shown to have reduced toxicity, both alone and in combination with a PARP inhibitor^20^.

Much like with other cancer therapies, identifying the subsets of tumors that respond well or are resistant to the drug will become paramount for efficient use of ATRi in the clinic. Genomewide screens are an effective technique to identify biomarkers of drug resistance and sensitivity. Recently reported genome-wide CRISPR knockout screens identified genetic determinants of ATRi sensitivity ^21,22^. By using a relatively low ATRi dose, these screens were designed to identify genes that, when lost, caused sensitivity to AZD6738. In contrast, little is known about genes that cause resistance to ATR inhibitors when inactivated. Moreover, these screens only investigated one ATR inhibitor. Additionally, genome-wide identification of genes that alter the response to ATRi when overexpressed rather than suppressed, has not yet been addressed.

To comprehensively identify the genes regulating the resistance to ATR inhibitors, we employed a dual CRISPR screening approach wherein we investigated both loss and overexpression of the majority of genes in the human genome. We performed both knockout and activation screens in HeLa cancer cells. Additionally, we performed the activation screens in nontransformed, breast epithelial MCF10A cells. All screens were performed separately with two different ATR inhibitors, VE822 and AZD6738. This comprehensive approach allowed unbiased identification of genes that affect ATRi resistance.

## Results

### Genome-wide CRISPR knockout screens identify genes regulating the resistance to multiple ATRi

A dual genome-wide CRISPR knockout and activation screening approach was designed to identify genes involved in the response to multiple ATR inhibitors (Figure 1). First, to identify genes whose loss confers resistance to ATRi, the Brunello human CRISPR knockout lentiviral library was employed^23^. This library targets 19,114 genes with 76,441 unique guide RNAs, thus on average covering each gene with four different gRNAs. To maintain 250X library coverage, 20 million library-infected cells were treated with VE822 (1.5μM), AZD6738 (3.6μM), or DMSO control. In contrast to previously published screens which focused on ATRi sensitivity and thus used a relatively low dose, we chose these high ATRi dosages as we previously determined that they kill approximately 90% of cells over 108 hours of treatment, thus allowing us to specifically study resistance to the drugs. Surviving cells were collected and genomic DNA was extracted. The sgRNA sequences were PCR-amplified and identified by Illumina sequencing (Figure 1A, C).

**Figure 1.**
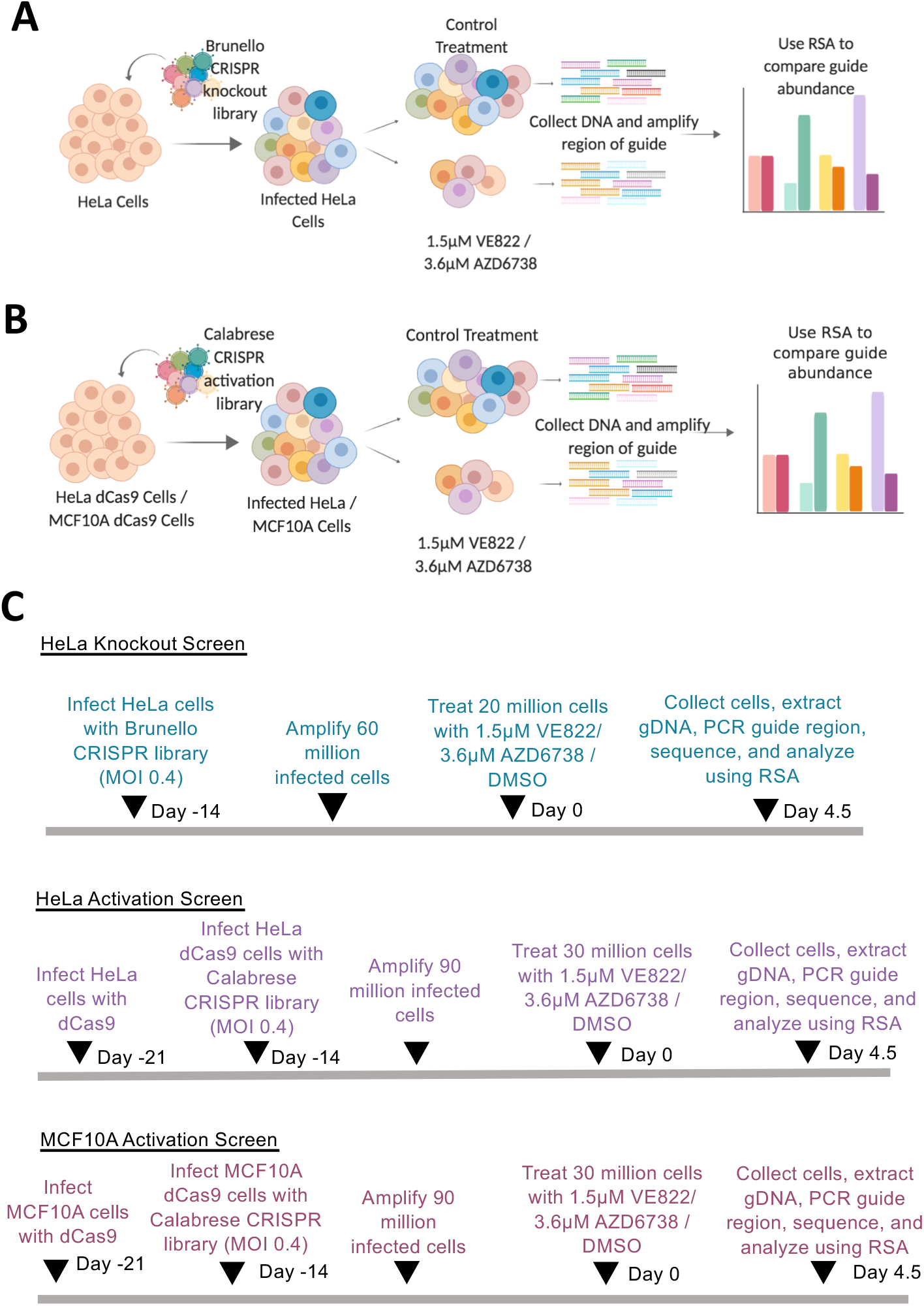
Overview of the dual CRISPR knockout and CRISPR activation screens to identify genes that regulate the resistance to the ATR inhibitors AZD6738 and VE822. (**A**) CRISPR knockout screens performed in HeLa cells using the Brunello CRISPR knockout library. (**B**) CRISPR activation screens performed in HeLa and MCF10A cells using the Calabrese CRISPR activation library. (**C**) Timelines of the knockout and activation CRSIPR screens.

Bioinformatic analyses using the redundant siRNA activity (RSA) algorithm^24^ was used to generate separate ranking lists of genes that were enriched in the VE822 and AZD6738 conditions compared to the control (Supplemental Table S1). This represents genes that, when inactivated, confer resistance to ATRi. Interestingly, there was large overlap between the VE822 and AZD6738 screen results, indicating common response mechanisms to the two different ATRi. Of the top 500 hits of each of the two ATRi screens, 155 were present in both of them (Figure 2A, Supplemental Table S2), which is much higher than the random probability (Figure 2B). Moreover, 7 genes were common within the top 40 hits of each ATRi screen. Biological pathway analysis of the top 500 hits of both screens revealed common biological processes, including DNA repair, translation, DNA replication, and sister chromatid cohesion (Figure 2C). Notably, multiple components of the cell cycle, cell migration, and DNA repair biological processes were found in both ATRi screens (Figure 2D).

**Figure 2.**
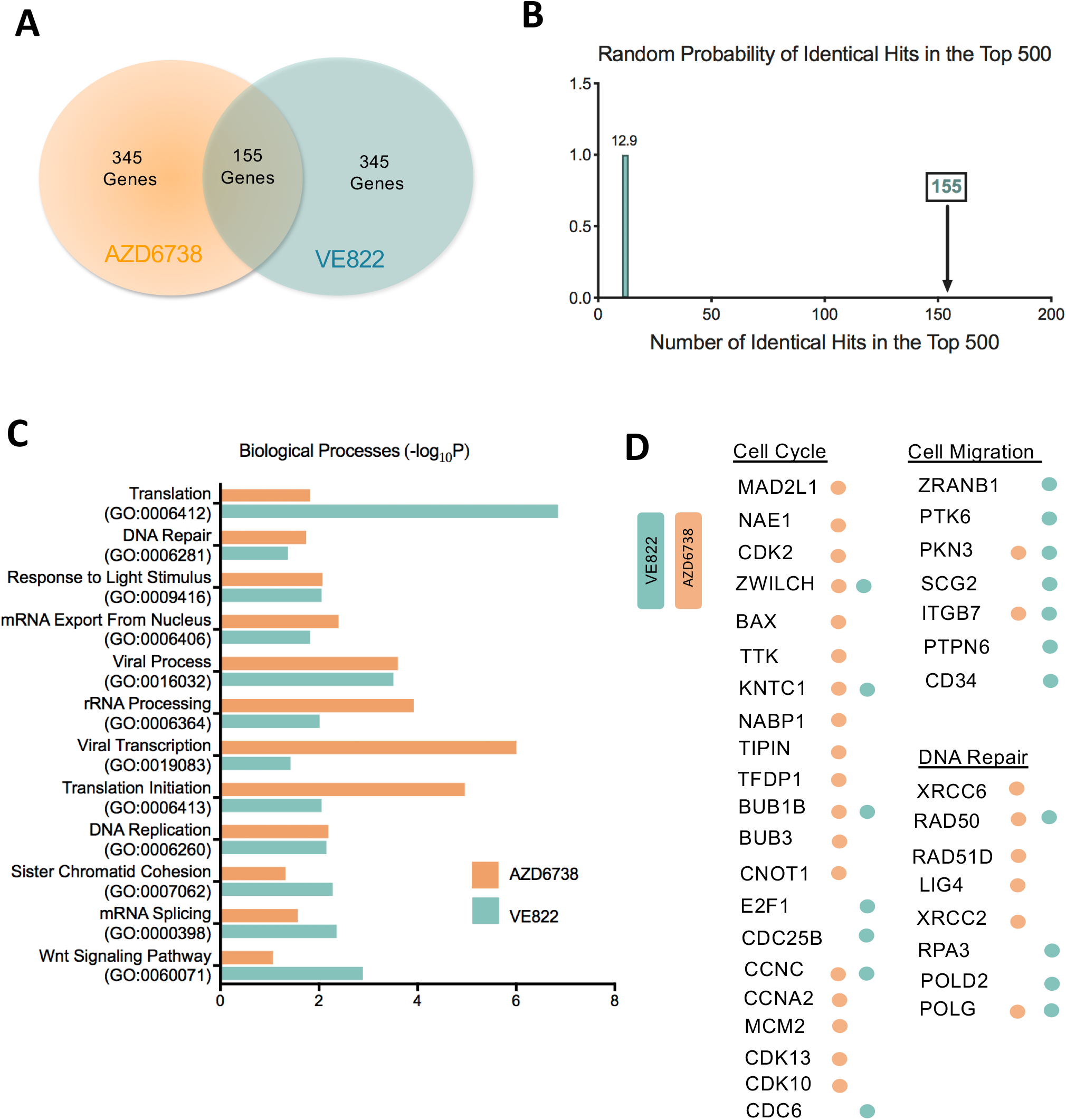
Pathway analysis reveals biological pathways involved in resistance to AZD6738 and VE822 from the CRISPR knockout screens. (**A**) Diagram showing the overlap of identical genes within the top 500 hits from both screens. (**B**) The number of common genes within the top 500 (namely 155) is much higher than the random probability of identical hits, which is 12.9. (**C**) Pathways that were significantly enriched in the top 500 hits from both ATRi screens using Gene Ontology analysis. (**D**) Genes in the top 500 hits of each ATRi screen that are involved in the indicated biological processes.

### Validation of hits from the knockout screen

Seven common genes were found within the top 40 genes in both ATRi resistance CRISPR knockout screens; these genes were: KNTC1, EEF1B2, LUC7L3, SOD2, MED12, RETSAT, and LIAS (Figure 3A, B). None of these genes were previously shown to induce ATRi resistance, to our knowledge. Thus, we sought to directly confirm that these seven genes alter the ATRi response. First, we tested these genes in HeLa cells, in which the screen was originally performed. We employed siRNA to knockdown each of these genes. The knockdown efficiency was confirmed by western blot for siSOD2, siMED12, siLUC7L3, and siEEF1B2, for which antibodies are available (Supplemental Figure S1). We performed clonogenic assays by incubating siRNA-treated HeLa cells with 1μM AZD6738 or 0.5μM VE822. After three days the media was replaced and colonies were allowed to grow for two weeks. HeLa cells with knockdown of each of the seven hits presented more colonies compared to control, indicating that they are more resistant to ATR inhibitors (Figure 3C). Moreover, we also validated the seven top hits by measuring cellular proliferation. HeLa cells were treated with siRNA and incubated for three days with 0.5μM VE822, 1μM VE822, 0.5μM AZD6738, or 1μM AZD6738. Cellular proliferation was determined using the CellTiterGlo reagent. Knockdown of each of the 7 top hits resulted in increased cell survival after ATR inhibitor treatment compared to control (Figure 3D). These findings show that all seven hits investigated confer resistance to both ATRi tested, thus validating our screen results.

**Figure 3.**
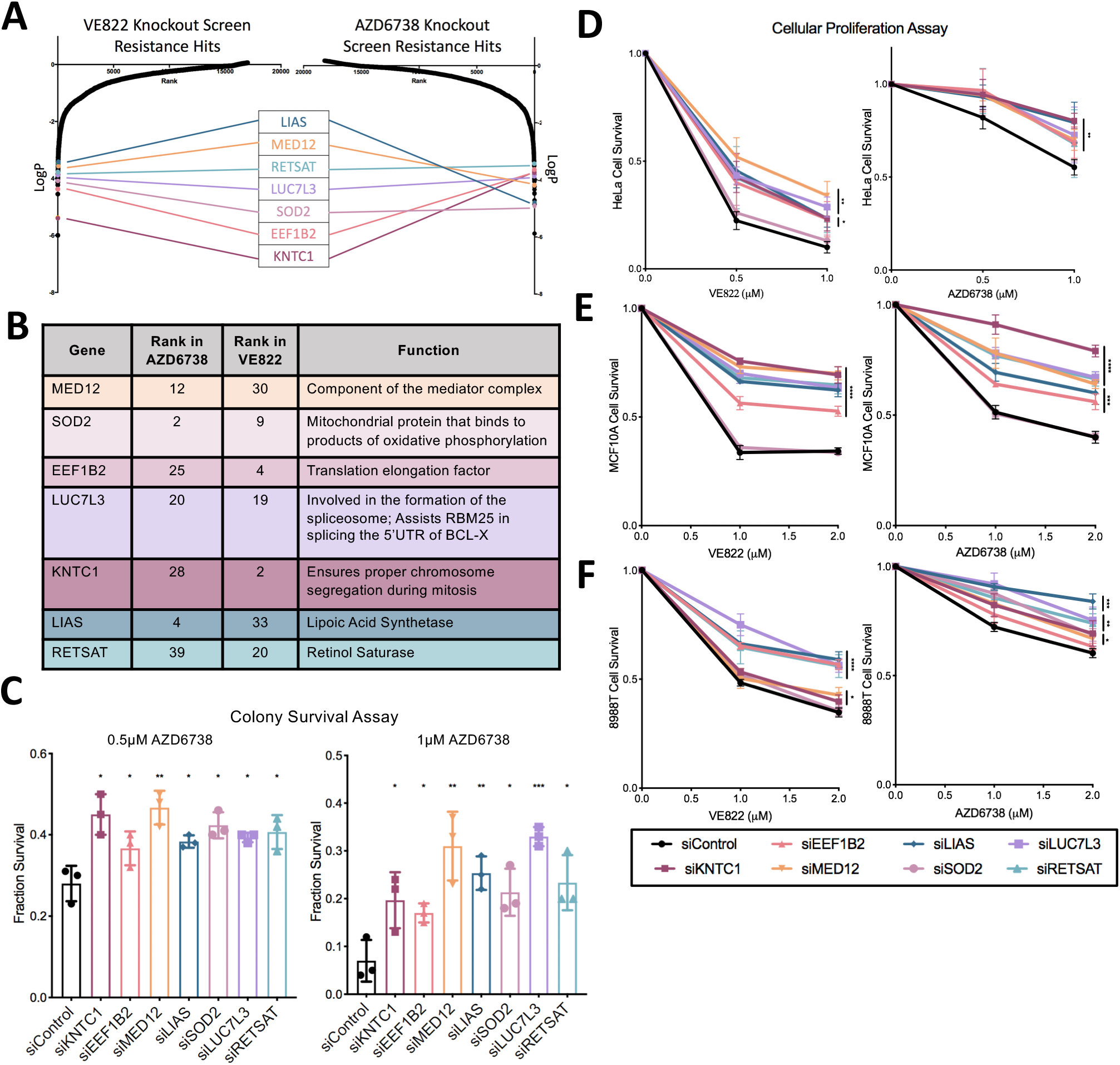
The top seven common hits were confirmed to cause resistance to ATRi when depleted in three different cell lines. (**A**) Scatterplot showing the results of the ATRi knockout screens. Each gene targeted by the library was ranked according to P-values calculated using RSA analysis. The P-values are based on the fold change of the guides targeting each gene between the ATRi- and DMSO-treated conditions. There were seven identical hits in the top 40 hits of both ATRi knockout screens. (**B**) The seven top common hits have diverse biological functions. (**C**) Knockdown of the top common gene hits in HeLa cells results in resistance to ATRi in a colony survival assay. The average of three experiments is shown, with error bars representing standard deviations. Asterisks indicate statistical significance for each hit compared to control. (**D**) Knockdown of the top common gene hits in HeLa cells results in resistance to ATRi in a cellular proliferation assay. The average of three experiments is shown, with error bars representing standard deviations. Asterisks indicate statistical significance for each hit compared to control. (**E, F**) Knockdown of the top common gene hits also results in resistance to ATRi in MCF10A cells (**E**) and 8988T cells (**F**) in cellular proliferation assays. The average of three experiments is shown, with error bars representing standard deviations. Asterisks indicate statistical significance for each hit compared to the control.

We next sought to investigate if these findings are restricted to HeLa cells, or in fact these hits also regulate ATRi resistance in other cell lines. To address this, we repeated the cellular proliferation experiments in MCF10A normal breast epithelial and 8988T pancreatic cancer cell lines. Both of these cell lines were slightly less sensitive to ATR inhibitors, so higher concentrations were used. Cells treated with 1μM VE822, 2μM VE822, 1μM AZD6738, and 2μM AZD6738 were analyzed for cellular proliferation after 3 days. In MCF10A cells, knockdown of all hits with the exception of SOD2 resulted in resistance to either ATR inhibitor (Figure 3E). In 8988T cells, knockdown of each of the 7 hits resulted in significant ATRi resistance (Figure 3F). These findings indicate that the genes identified by screening HeLa cells control ATRi resistance across multiple cell lines.

### Loss of the top hits does not restore ATR catalytic activity

One potential mechanism of ATRi resistance is the restoration of ATR catalytic activity in the presence of ATRi. Chk1 is the main component of the ATR signaling cascade that is activated by DNA damage^25^. Once ATR is activated, its kinase activity phosphorylates Chk1 leading to all of the downstream effects. As a result, Chk1 phosphorylation at Serine 317 and Serine 345 represent markers of ATR cascade activation and activity. Thus, we knocked down the top hits and analyzed the levels of phosphorylated Chk1 by western blot, under no treatment conditions, hydroxyurea treatment, ATRi treatment, and a combination of hydroxyurea and ATRi. Hydroxyurea depletes the cellular dNTP pools, thereby stimulating the replication stress response and activating ATR. We found that after knockdown of our top hits, there was no difference in Chk1 phosphorylation by any of the hits (Supplemental Figure S2). This indicates that these genes do not act by restoring ATR activity in the presence of ATRi.

### Loss of LUC7L3 causes resistance to ATRi by suppressing ATRi-induced apoptosis through splicing of BCL-X

Treatment of HeLa cells with VE822 for three days resulted in induction of apoptosis, as measured by detecting Annexin-V expression (Figure 4A). Thus, we sought to employ this assay as another readout of resistance to ATR inhibitors. Knockdown of each of the 7 top hits significantly reduced the amount of apoptosis induced by treatment with 0.5μM VE822 for three days (Figure 4B), further validating that their loss confers resistance to ATRi.

**Figure 4.**
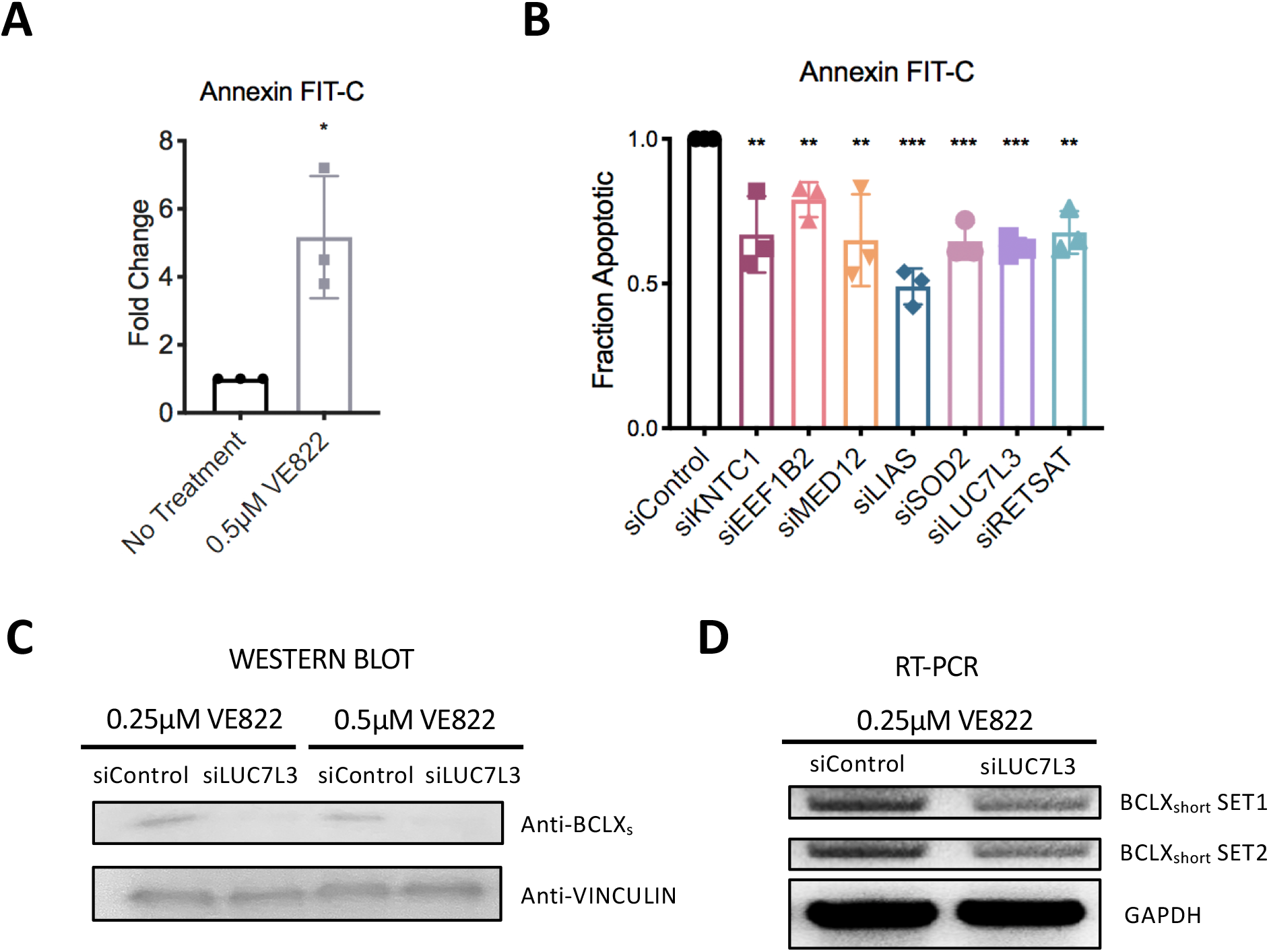
Regulation of ATRi-induced apoptosis by LUC7L3. (**A**) Treatment with ATR inhibitors results in apoptosis induction, as measured by the Annexin-V assay. The average of three experiments is shown, with error bars representing standard deviations. Asterisks indicate statistical significance. (**B**) Knockdown of the top 7 hit genes in HeLa cells causes a significant reduction in apoptosis after treatment with 0.5μM VE822 for 36 hours. First, data for each hit was normalized to their own untreated control, and then normalized to siControl. The average of three experiments is shown, with error bars representing standard deviations. Asterisks indicate statistical significance for each hit compared to control. (**C**) Western blot showing a decrease in the amount of the BCLX_short_ isoform in HeLa cells after knockdown of LUC7L3 and ATRi treatment. (**D**) Reverse transcriptase PCR showing a decrease in BCLX_short_ mRNA after knockdown of LUC7L3 and ATRi treatment in HeLa cells.

One of the top hits, namely LUC7L3, was previously identified as an interactor of RBM25, an mRNA splicing factor which regulates the splicing of the apoptosis regulator BCL-X^26^. The long isoform of BCL-X is anti-apoptotic, while the short isoform is pro-apoptotic^27^. Loss of RBM25 causes a shift in the splicing of BCL-X from the short isoform to the long isoform. Thus, we hypothesized that loss of LUC7L3 may cause a decrease in the pro-apoptotic BCL-X_short_, which could help explain the resistance to ATRi-induced apoptosis. To test this, we knocked-down LUC7L3 in HeLa cells and treated these cells with 0.25μM or 0.5μM VE822 for 24 hours. We then collected the cells and investigated BCL-X_short_ levels at both the protein and mRNA levels. Western blots using an antibody specific to BCL-X_short_ showed a reduction in the levels of this isoform in LUC7L3-depleted cells (Figure 4C). Moreover, using reverse transcriptase PCR we observed a decrease in BCLX_short_ mRNA levels in the cells treated with siRNA targeting LUC7L3 compared to control (Figure 4D). These results indicate that the loss of LUC7L3 causes a decrease in the pro-apoptotic isoform of BCL-X, therefore potentially explaining the resistance to ATR inhibitor treatment.

### Loss of MED12 and LIAS stabilizes the replication fork in response to ATRi

A major role of ATR is to stabilize replication forks upon genotoxic insults^28^. ATR inhibitors have been shown to cause a decrease in replication tract length^15^. This replication fork slowing is detrimental to the cell and may eventually result in replication deficiency and cell cycle arrest. We sought to investigate if any of the hits described above regulate ATRi resistance by suppressing the impact of ATRi on fork slowing. For this, we employed the DNA fiber assay. We incubated cells consecutively with two thymidine analogs to detect active DNA replication forks. IdU was added first for 30 minutes followed by CldU for 30 minutes. VE822 was added concomitantly with CldU. With increasing amounts of ATRi (2μM VE822, 4μM VE822, or 6μM VE822), the CldU replication tract was significantly decreased (Figure 5A), thereby confirming that ATR inhibition causes a decrease in replication fork progression.

**Figure 5.**
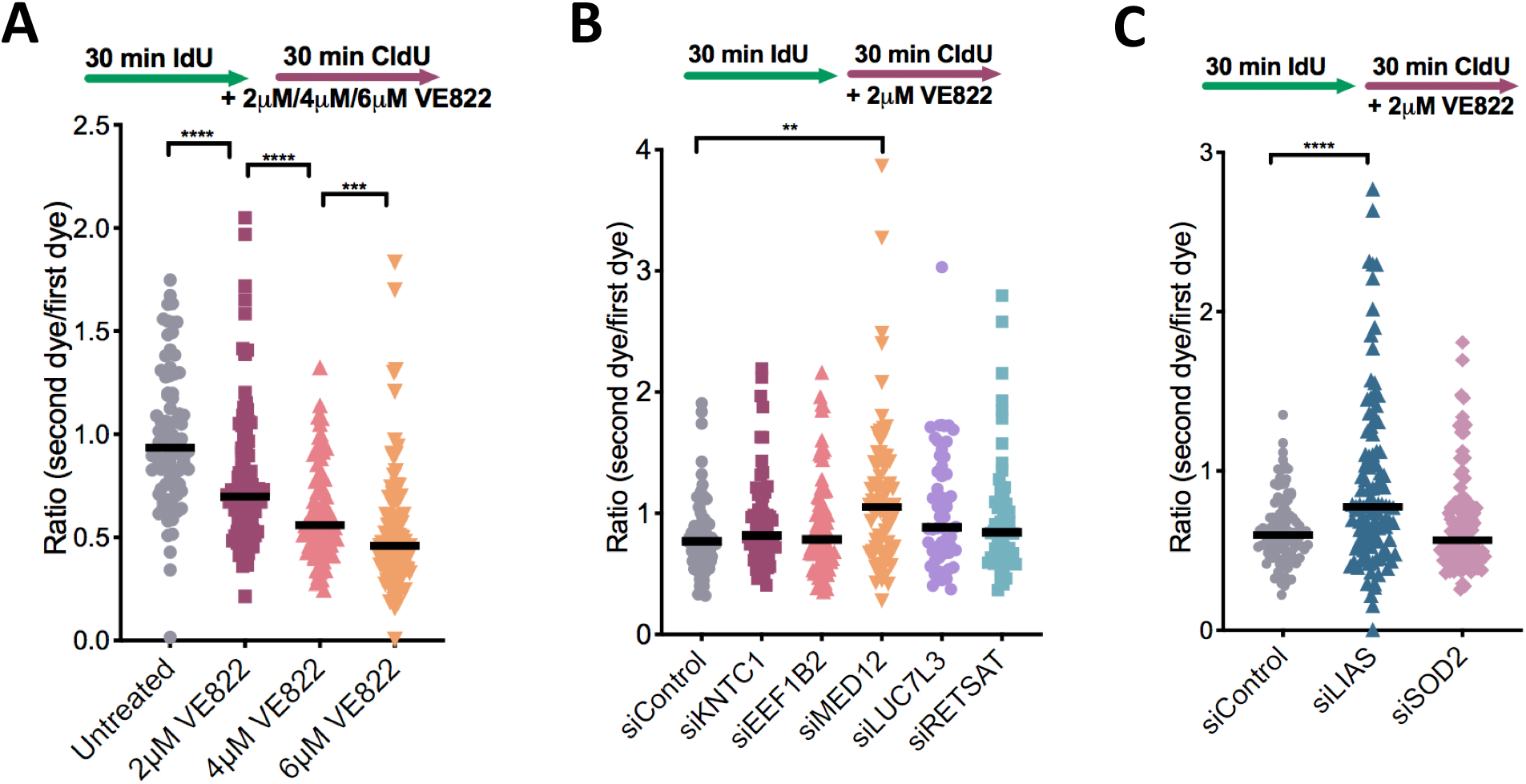
Knockdown of MED12 and LIAS promotes replication fork stability in the presence of ATRi. (**A**) The addition of increasing concentration of ATRi in HeLa cells causes a significant decrease in replication tract length, as measured by the ratio of the CldU tract length (without ATRi) to the IdU tract length (with ATRi). (**B, C**) Upon treatment with 2μM VE822 concomitant with CldU, knockdown of MED12 (**B**) or LIAS (**C**) showed a restoration of normal fork speed, as measured by the ratio of the CldU tract length (without ATRi) to the IdU tract length (with ATRi). The other hits do not affect fork slowing induced by ATRi. In all panels, the mean values are indicated for each sample, and the asterisks indicate statistical significance. At least 100 fibers were quantified.

Next, we investigated the impact of the top hits described above, by performing the DNA fiber combing assay with cells depleted of each of these genes by siRNA. Cells were treated with IdU for 30 minutes followed by co-treatment with CldU and 2μM VE822 for 30 minutes. The effect of loss of the top hits was investigated by calculating the ratio of CldU tract length to IdU tract length. Strikingly, knockdowns of MED12 and LIAS significantly increased the CldU-to-IdU ratio, indicating that fork slowing in the presence of ATRi is suppressed (Figure 5B, C). The other hits did not affect the CldU-to-IdU ratio. Therefore, the ATRi resistance observed upon loss of MED12 and LIAS may involve restoration of replication fork speed in the presence of ATR inhibitors.

### Genome-wide CRISPR activation screens identify genes whose overexpression causes resistance to multiple ATRi

To evaluate genes whose overexpression leads to resistance to ATR inhibitors, we performed CRISPR activation screens in HeLa cells as well as MCF10A cells. The Calabrese CRISPR activation library was employed for these experiments^29^. This library targets 18,885 genes with 56,762 sgRNAs, for an average of 3 guides per gene. First, HeLa and MCF10A cells were infected with a lentivirus containing the dCas9 construct necessary for transcriptional activation (Supplemental Figure S3A). Expression of dCas9 was confirmed by Western blot (Supplemental Figure S3B). These cells were then infected with the activation library. Next, 30 million library-infected cells (for 500X library coverage) were treated with ATRi using the same conditions as for the knockout screens (Figure 1). Surviving cells were collected and genomic DNA was extracted. The sgRNA sequences were PCR-amplified and identified by Illumina sequencing (Figure 1B, C). Using the RSA algorithm, we generated separate lists of genes that were enriched in the VE822 and AZD6738 conditions compared to the control (Supplemental Table S3). This represents genes that, when overexpressed, confer resistance to ATRi. Similar to the knockout screen, there was large overlap between the VE822 and AZD6738 conditions. Within the top 500 genes, 99 genes were common for the two ATRi screens in HeLa cells (Figure 6A, Supplemental Table S4), and 115 genes were common in MCF10A screens (Figure 6B, Supplemental Table S4). The number of common hits is much higher than expected from a random distribution (Figure 6C). Moreover, within the top 40 genes, 7 hits were common in HeLa cells (Figure 6D) and 6 were common in MCF10A cells (Figure 6E). Importantly, all seven top hits from the HeLa knockout screen, were ranked towards the bottom in the HeLa overexpression screens (Figure 7A), indicating that their overexpression promotes ATRi sensitivity and thus further highlighting the relevance of our screening strategy.

**Figure 6.**
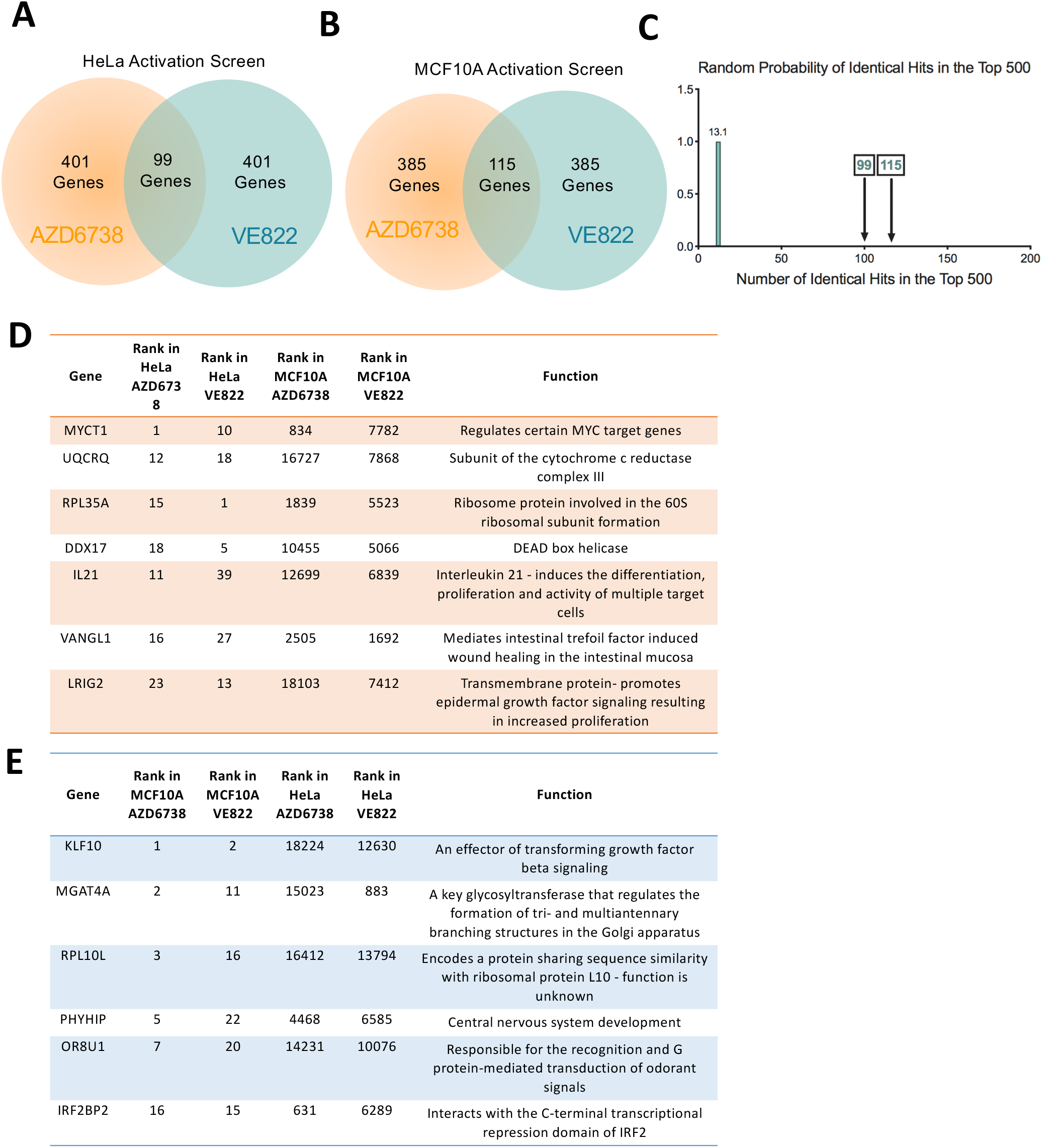
CRISPR activation screens in HeLa and MCF10A cells to identify genes that result in resistance to multiple ATRi when overexpressed. (**A**) Diagram showing the overlap of identical genes within the top 500 hits from both ATRi activation screens in HeLa cells. (**B**) Diagram showing the overlap of identical genes within the top 500 hits from both ATRi activation screens in MCF10A cells. (**C**) The number of common genes within the top 500 (namely 99 for the HeLa screens and 115 for the MCF10A screens) is much higher than the random probability of identical hits. (**D, E**) Tables listing the common genes among top 40 hits in each of the ATRi CRISPR activation screens in HeLa (**D**) and MCF10A (**E**) cells.

**Figure 7.**
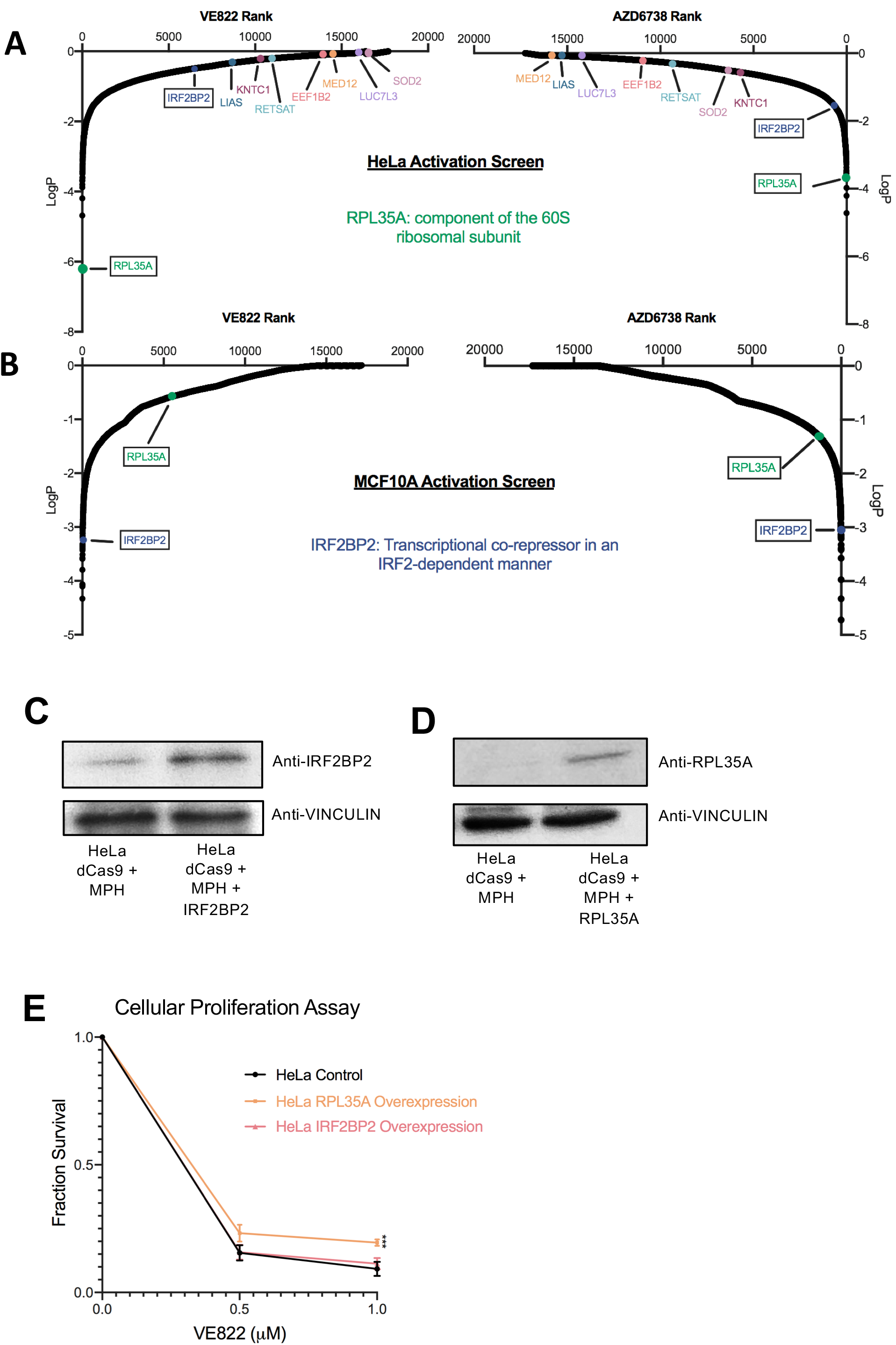
Validation of the CRISPR activation screens for ATRi resistance. (**A**) Scatterplot showing the results of the ATRi activation screens in HeLa cells. Each gene targeted by the library was ranked according to P-values calculated using RSA analysis. The P-values are based on the fold change of the guides targeting each gene between the ATRi- and DMSO-treated conditions. The location of RPA35A and IRF2BP2 is shown (in black boxes). RPL35A is a top hit in both ATR inhibitors HeLa screens, whereas IRF2BP2 is not. Also shown is the location of the seven top hits from the HeLa knockout screen described above. (**B**) Scatterplot showing the results of the ATRi activation screens in MCF10A cells. IRF2BP2 is a top hit in both ATRi screens, whereas RPL35A is not. (**C**) Western blot showing the overexpression of IRF2BP2 in HeLa cells. (**D**) Western blot showing the overexpression of RPL35A in HeLa cells. (**E**) Overexpression of RPL35A in HeLa cells causes resistance to VE822 in a cellular proliferation assay, while overexpression of IRF2BP2 does not. The average of three experiments is shown, with error bars representing standard deviations. Asterisks indicate statistical significance compared to control.

Interestingly, when comparing between the HeLa and MCF10A screens, there was no overlap of top hits (Supplemental Table S3). This surprising finding perhaps reflects the inherent differences between these cell lines, as one is cancer-derived and p53 deficient, while the other is non-transformed with wild-type p53 function. To confirm these findings, we chose to validate two hits that were common to both ATR inhibitors, but cell line specific. RPL35A was a top hit for both ATRi screens in HeLa cells, but was not a top hit in the MCF10A screens (Figure 7A, B). On the other hand, IRF2BP2 was a top hit for both ATRi screens in the MCF10A cells, but not in the HeLa screens (Figure 7A, B). We overexpressed these two genes in the HeLa dCas9 cells by CRISPR activation using the MS2-P65-HSF1 (MPH) activator complex, to induce transcription. Western blots confirmed the overexpression of both IRF2BP2 (Figure 7C) and RPL35A (Figure 7D). Cellular proliferation of the overexpression cell lines was analyzed upon incubation with VE822 for three days. RPL35A overexpression resulted in ATRi resistance compared to control cells, while IRF2BP2 overexpression did not show any difference to control cells (Figure 7E). These results are in line with the screen data that identified RPL35A as a top hit in HeLa cells but not in MCF10A cells (Figure 7A, B), thus validating the activation screens. These findings suggest that different mechanisms may regulate ATRi resistance in HeLa compared to MCF10A cells.

## Discussion

Detailed information on the genetic make-up of tumors will help to better treat patients on a personalized basis. Identification of markers that lead to resistance to ATRi is critical to advance the use of ATRi in cancer therapy. With the emerging use of ATRi in clinical trials, there has been renewed interest in determining predictors to ATRi response. Moreover, beyond the treatment of patients, the effects of ATRi on cell biology and pathways that regulate responses to ATRi are still not well understood.

Here, we present dual-genome wide CRISPR knockout and activation screens to identify predictors of ATRi resistance in HeLa and MCF10A cells. Specifically, two of the main ATRi that are currently in clinical trials, VE822 and AZD6738 were investigated. Our CRISPR screening strategy provides an unbiased approach to identify genes that, when altered, cause resistance to ATRi. First, we performed a CRISPR knockout screen in HeLa cells to identify genes that, when knocked out, caused resistance to VE822 and AZD6738. When comparing the results for the two ATRi screens, we observed a significant overlap of the top hits. 155 of the top 500 genes were common between the two ATRi, as were 7 of the top 40. This high number of identical hits, much higher than expected from a random distribution, validates our screening strategy and shows that common pathways are involved in the response to different ATRi.

We validated that loss of these seven top-ranked common gene hits causes resistance to both VE822 and AZD6738, in three different cell lines using siRNA knockdown. Mechanistically, we show that none of these cases involve restoration of ATR activity in the presence of the inhibitors. Instead, we identified two different mechanisms through which some of these genes regulate ATRi resistance. We found that LUC7L3 regulates the splicing of the BCL-X apoptosis regulator. Upon loss of LUC7L3, the level of the pro-apoptotic isoform of BCL-X decreases, resulting in reduced ability to undergo apoptosis, potentially explaining the ATRi inhibitor resistance of these cells.

Another mechanism of resistance we uncovered involves restoration of replication fork protection. It was previously shown that ATR inhibition causes a decrease in fork progression and an increase in origin firing^28^. Under normal conditions, ATR is responsible for suppression of local origin firing, therefore when ATR is inhibited, origin firing increases^30^. As there is an inverse correlation between DNA tract length and number of origins firing, the more origins that fire, the slower the fork progression rate is^31^. However, in the case of ATRi it is still unknown whether replication slowing causes more origins to fire, or if more origins firing cause the replication forks to slow^32^. We found that loss of either MED12 or LIAS causes a restoration of fork stability as the decrease in fork progression upon ATRi treatment is not seen under these conditions. This result suggests that mechanisms countering fork slowing or origin firing in the presence of ATRi, may in fact promote cellular viability under these conditions.

Finally, we completed the first (to our knowledge) genome-wide CRISPR activation screen to identify genes that cause resistance to ATRi when overexpressed. This activation screen was performed in both HeLa and MCF10A cells. Validating the screening strategy, top hits from the knockout screen ranked very low on the list of hits for the activation screens. Similar to the HeLa knockout screen, there was a high number of identical top hits between the VE822 and AZD6738 screens for each cell line. However, there was almost no overlap between the two cell lines. We confirmed these surprising results by showing that overexpression of RPL35A, a top hit from the HeLa screens, causes ATRi resistance in HeLa cells, but overexpression of IRF2BP2, a top hit from the MCF10A screens, does not cause resistance in HeLa cells. These findings reflect the inherent differences between the two cell lines, as HeLa cells are tumor-derived, whereas MCF10A cells are non-transformed breast epithelial cells. These results suggest that biomarkers of gene overexpression specific to tumor cells may be used to create treatment plans with fewer side effects to the rest of the body.

Our studies employed a unique combination of genome-wide CRISPR-based screening approaches to comprehensively identify genes that, when altered, cause resistance to ATRi. We analyzed two separate ATRi using both knockout and activation screens, in both cancer cells and non-transformed cells. Our findings could provide biomarkers to ultimately help create effective treatment plan for cancer therapy with ATR inhibitors.

## Materials and Methods

### Cell Culture

HeLa and 8988T cells were grown in Dulbecco’s modified Eagle’s medium (DMEM) supplemented with 10% fetal calf serum and 1% Pen/Strep. MCF10A cells were grown in DMEM/F12 supplemented with 5% fetal calf serum, 1% Pen/Strep, 20ng/mL hEGF, 0.5 mg/mL Hydrocortisone, 100 ng/mL Cholera Toxin, and 10 μg/mL Insulin.

Gene knockdown was performed using Lipofectamine RNAiMAX transfection reagent. Cells were treated with siRNA for two consecutive days. The following SilencerSelect oligonucleotides (ThermoFisher) were used for gene knockdown: KNTC1 (ID: s18776); LUC7L3 (ID: s226748); SOD2 (ID: s13267); LIAS (ID: s223178); EEF1B2 (ID: s194388); RETSAT (ID: s29671); MED12 (ID: s19362). AllStars Negative Control siRNA (Qiagen 1027281) was used as control.

HeLa cells overexpressing RPL35A and IRF2BP2 were created by consecutive rounds of transduction and selection to induce transcriptional activation. Cells were first transduced with the dCas9 lentiviral construct (Addgene 61425-LV) and selected with 3μg/ml blasticidin. The resulting HeLa-dCas9 cells were then transduced with the lentiviral construct for the MS2-P65-HSF1 (MPH) activator complex (Addgene 61426-LVC) and selected with 0.5 mg/ml hygromycin. Finally, HeLa-dCas9-MPH cells were transduced with lentivirus constructs containing the following guide sequences: CGGTGGCGGCCGCGTCCCGG for IRF2BP2, and CAGTGCGAAGCCGATTTCCG for RPL35A (Sigma Custom CRISPR in lentiviral backbone LV06).

### Protein Techniques

Cell extracts and western blots were performed as previously described^33^. Antibodies used were: MED12 (Santa Cruz Biotechnology sc-515695), SOD2 (Santa Cruz Biotechnology sc-133254), LUC7L3 (ProteinTech 145041AP), EEF1B2 (ProteinTech 104831AP), BCLX_short_ (Invitrogen PA5-78864), Vinculin (Santa Cruz Biotechnology sc-73614), GAPDH (Santa Cruz Biotechnology sc-47724), CAS9 (BioLegend 844302), pChk1-S317(Cell Signaling 2344S), pChk1-S345(Cell Signaling 2341S), IRF2BP2 (ProteinTech 188471AP), RPL35A (Bethyl, 501569555).

### CRISPR Screens

For CRISPR knockout screens, the Brunello Human CRISPR knockout pooled lentiviral library (Addgene 73179) was used^23^. This library is comprised of 76,411 gRNAs that target 19,114 genes. Fifty million HeLa cells were infected with this library at a multiplicity of infection (MOI) of 0.4 to achieve 250X coverage and selected for 4 days with 0.6 μg/mL puromycin. For ATRi resistance screens, 20 million library-infected cells (to maintain 250x coverage) were used for each drug condition: DMSO (vehicle control), 1.5μM VE822 (Selleck S7102), and 3.6μM AZD6738 (Selleck S7693). Cells were treated for 96 hours and then collected. Compared to control cells, survival of ATRi-treated cells was 8% (for VE822) and 10% (for AZD6738) respectively.

For CRISPR activation screens, the Calabrese Human CRISPR activation pooled library, targeting 18,885 genes with 56,762 gRNAs, was used (Set A, AddGene 92379)^29^. First, wild-type HeLa and MCF10A cells were infected with the dCas9 lentiviral construct (Addgene 61425-LV) and selected with 3μg/ml blasticidin for 5 days. The presence of the dCas9 was confirmed by western blot using a Cas9 antibody. Next, 75 million HeLa-dCas9 and MCF10A-dCas9 cells were infected with this library at a MOI of 0.4 to achieve 500x coverage and selected for 96 hours with 0.6 μg/mL puromycin. For ATRi resistance screens, 30 million cells were used for each drug treatment condition to maintain 500x coverage of the library. Library-infected HeLa-dCas9 and MCF10A-dCas9 cells were treated with DMSO (vehicle control), 1.5μM VE822, or 3.6μM AZD6738 for 96 hours. For HeLa cells, survival of ATRi treated cells was 9% (VE822) and 11% (AZD6738), respectively. For MCF10A cells, survival was 15% (VE822) and 13% (AZD6738), respectively.

### Sequencing and analysis of CRISPR screens

Genomic DNA was isolated using the DNeasy Blood and Tissue Kit (Qiagen 69504) per the manufacturer’s instructions. gRNAs were identified in our populations using PCR primers with Illumina adapters. Genomic DNA from a number of cells corresponding to the equivalent of 250-fold library coverage was used as template for PCR (20 million cells for knockout screen, 15 million cells for activation screen). 10μg of gDNA was used in each PCR reaction along with 20μl 5X HiFi Reaction Buffer, 4μl of P5 primer, 4μl of P7 primer, 3μl of Radiant HiFi Ultra Polymerase (Stellar Scientific), and water. The P5 and P7 primers were determined using the user guide provided with the CRISPR libraries (https://media.addgene.org/cms/filer_public/61/16/611619f4-0926-4a07-b5c7-e286a8ecf7f5/broadgpp-sequencing-protocol.pdf). The PCR cycled as follows: 98°C for 2min before cycling, then 98°C for 10sec, 60°C for 15sec, and 72°C for 45sec, for 30 cycles, and finally 72°C for 5min. After PCR purification, the final product was Sanger sequenced to confirm that the guide region is present, followed by qPCR to determine the exact amount of PCR product present. The purified PCR product was then sequenced with Illumina HiSeq 2500 single read for 50 cycles, targeting 10M reads.

Next, the sequencing results were analyzed bioinformatically. First, the sgRNA representation was analyzed using the custom python script provided (count_spacers.py)^34^. The difference between the number of guides present in each ATRi condition compared to control condition was then determined. Specifically, one read count was added to each sgRNA, and then the treatment reads were normalized to no treatment. Finally, the values found were used as input in the Redundant siRNA Activity (RSA) algorithm^35^. For RSA, the Bonferroni option was used and guides that were 2-fold enriched in treatment compared to no treatment were considered hits. This analysis method allows for the identification of genes that are upregulated in one population (VE822 or AZD6738) compared to control. Hits are determined by the amount of gRNA sequences present in the population and the number of guides per gene present. Furthermore, p-values are determined by the RSA algorithm for the genes that are most enriched in the test populations compared to the control. VE822 and AZD6738 top hits can then be compared. Biological pathway analysis of the top hits was performed using Gene Ontology^36,37^.

### Drug Sensitivity Assays

Cellular proliferation was measured using the CellTiterGlo reagent (Promega) according to the manufacturer’s instructions. After 2 days of siRNA treatment, 1500 cells per condition were plated into 96-well plates and treated with the indicated doses of ATRi for three days. CellTiterGlo reagent was added for 10 minutes before the luminescence was read on a plate reader. For colony survival assays, 500 siRNA treated cells were plated into 6-well plates and treated with the indicated doses of ATRi. After 3 days of treatment, media was replaced. After two weeks, cells were fixed and colonies were stained with crystal violet. Apoptosis was quantified using the FITC Annexin V kit (Biolegend 640906) according to the manufacturer’s instructions.

### DNA Fiber Combing Assay

HeLa cells were treated with siRNA as indicated, then incubated with 100μM IdU in DMEM for 30 minutes. Cells were then washed three times with PBS and incubated with 100μM CldU and VE822 as indicated, in DMEM for 30 minutes. Next, cells were collected and processed using the FiberPrep kit (Genomic Vision EXT-001) according to the manufacturer’s instructions. DNA molecules were stretched onto coverslips (Genomic Vision COV-002-RUO) using the FiberComb Molecular Combing instrument (Genomic Vision MCS-001). The slides were then stained with antibodies detecting CldU (Abcam 6236), IdU (BD 347580), and DNA (Millipore Sigma MAD3034). Next, slides were incubated with secondary Cy3, Cy5, or BV480-conjugated antibodies (Abcam 6946, Abcam 6565, and BD Biosciences 564879). Finally, the cells were mounted onto coverslips and imaged using confocal microscopy (Leica SP5).

### Reverse Transcriptase PCR (RT-PCR)

To detect the long and short isoforms of BCL-X mRNA, RT-PCR was used. RNA was extracted using TRIzol reagent (Life Tech) according to the manufacturer’s instructions. RNA was then converted to cDNA using the RevertAid Reverse Transcriptase Kit (Thermo Fisher Scientific) with oligo-dT primers. Next, PCR was performed with the following primers: GAPDH (for: TGCACCACCAACTGCTTAGC; rev: TCAGCTCAGGGATGACCTTG), BCLX-s/l set 1 (for: AGTAAAGCAAGCGCTGAGGGAG; rev: ACTGAAGAGTGAGCCCAGCAGA)^38^, BCLX-s/l set 2 (for: GAGGCAGGCGACGAGTTTGAA; rev: TGGGAGGGTAGAGTGGATGGT)^39^. The PCR cycled as follows: 95°C for 2min to start, followed by 95°C for 30sec, 45°C for 1min, and 68°C for 1min (cycled 30 times), with a final 5-minute incubation at 68°C. The PCR product was then run on a 2% agarose gel and imaged with a ChemiDoc Gel Imager (Bio-Rad).

### Statistical Analysis

The Mann-Whitney statistical test was performed for the DNA fiber assay. The CellTiterGlo proliferation assays were analyzed with a 2-way ANOVA. For other assays, the t-test (two-tailed, unequal variance unless indicated) was performed. Statistical significance is indicated for each graph (ns = not significant, for P > 0.05; * for P ≤ 0.05; ** for P ≤ 0.01; *** for P ≤ 0.001;**** for P ≤ 0.0001).

## Acknowledgements

We would like to thank Dr. Hong-Gang Wang and Dr. Tom Spratt for materials and advice; and the following Penn State College of Medicine core facilities: Flow Cytometry, Genomic Analyses, and Imaging. This work was supported by: NIH R01ES026184 and R01GM134681 (to GLM) and 1F31CA243301-01 (to EMS).

## Legends to Supplemental Tables

**Supplemental Table S1 – Lists of all genes in the CRISPR knockout screens ranked by p-value (2 tabs).**

**Supplemental Table S2 – List of common hits from the knockout screen (1 tab).**

**Supplemental Table S3 – Lists of all genes in the CRISPR activation screens ranked by p-value (4 tabs).**

**Supplemental Table S4 – List of common hits from the activation screens (2 tabs).**

## Legends to Supplemental Figures

**Supplemental Figure S1. Validation of siRNA knockdown of the top hits.** Western blots showing knockdown of SOD2 **(A)**, MED12 **(B)**, LUC7L3 (**C**), and EEF1B2 (**D**) are presented.

**Supplemental Figure S2. Levels of pChk1 are not restored upon knockdown of the top hits and subsequent treatment with ATRi.** Western blots showing the levels of pCHK1 S317 (**A**) and pChk1 S345 (**B**) in HeLa cells after knockdown of the top hits followed by 24 hours of no treatment, hydroxyurea treatment, ATRi treatment, or hydroxyurea with ATRi treatment, are shown.

**Supplemental Figure S3. Overview of the CRISPR activation screen. (A)** Schematic representation of the CRISPR activation screen setup. The gRNA targets dCas9 to the promoter region of the gene of interest, along with multiple transcriptional activators to upregulate the transcription of the gene. (**B**) Western blots showing dCas9 expression in the cells used for the CRISPR activation screen and HeLa-dCas9 overexpression cell lines.

