## Supplementary figures and images for "Dual genome-wide CRISPR knockout and CRISPR activation screens identify common mechanisms that regulate the resistance to multiple ATR inhibitors"

### Supplemental Figures S1-S3

#### Supplemental Figure S1

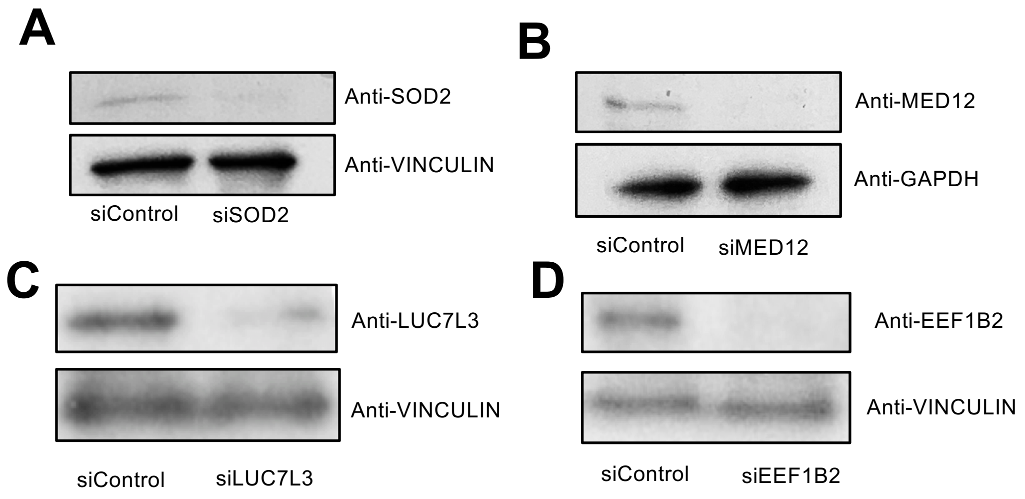

#### Supplemental Figure S2

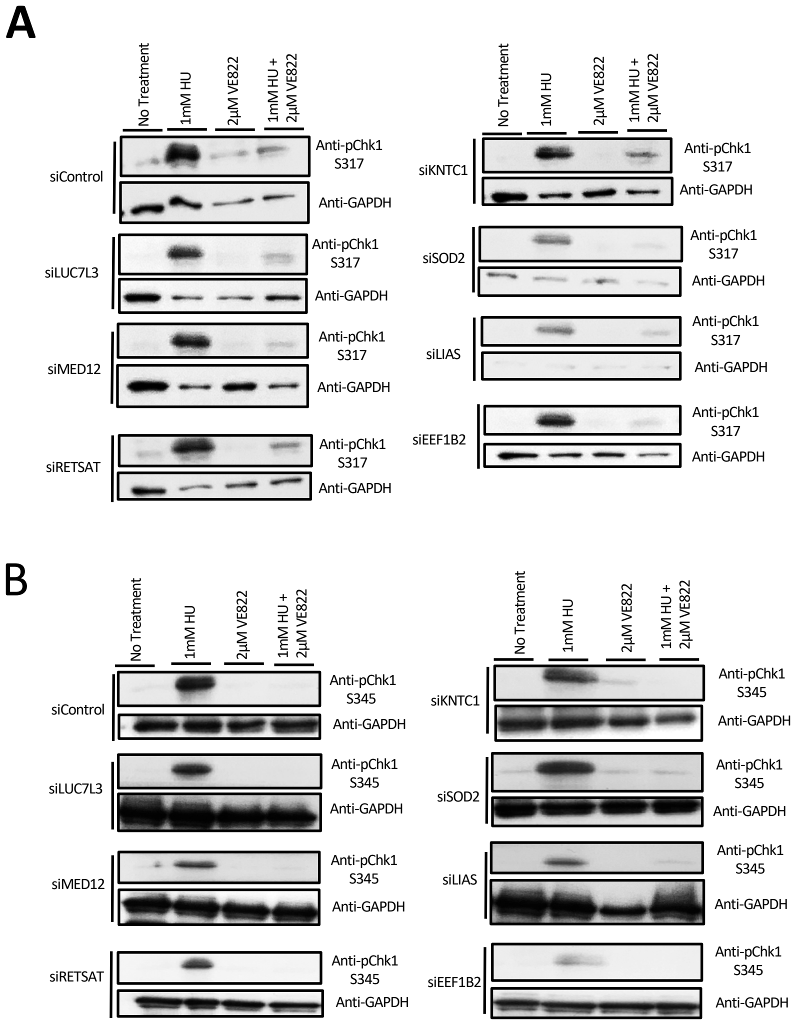

#### Supplemental Figure S3

**A**

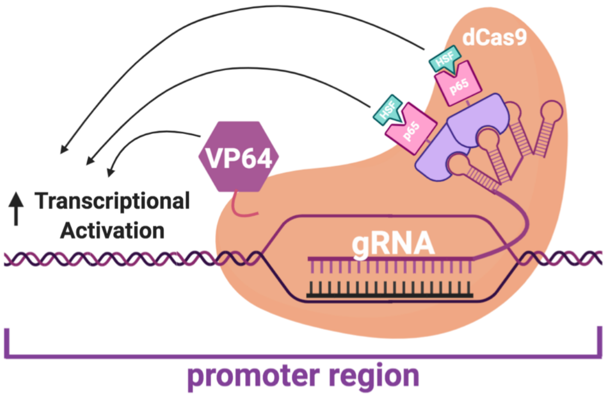


**B**

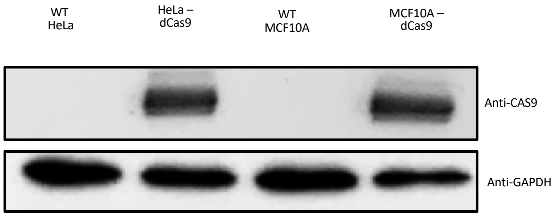
